# Newly developed MAGIC population allows identification of strong associations and candidate genes for anthocyanin pigmentation in eggplant

**DOI:** 10.1101/2021.09.10.459758

**Authors:** Giulio Mangino, Andrea Arrones, Mariola Plazas, Torsten Pook, Jaime Prohens, Pietro Gramazio, Santiago Vilanova

## Abstract

MAGIC populations facilitate the genetic dissection of complex quantitative traits in plants and are valuable breeding materials. We report the development of the first eggplant MAGIC population (S3MEGGIC; 8-way), constituted by 420 S3 individuals developed from the intercrossing of seven cultivated eggplant (*Solanum melongena*) and one wild relative (*S. incanum*) parents. The S3MEGGIC recombinant population was genotyped with the eggplant 5k probes SPET platform and phenotyped for anthocyanins presence in vegetative plant tissues (PA) and fruit epidermis (FA), and for the light-sensitive anthocyanic pigmentation under the calyx (PUC). The 7,724 filtered high-confidence SNPs confirmed a low residual heterozygosity (6.87%) and a lack of genetic structure in the S3MEGGIC population, including no differentiation among subpopulations carrying cultivated or wild cytoplasm. Inference of haplotype blocks of the nuclear genome revealed an unbalanced representation of founder genomes, suggesting cryptic selection in favour or against specific parental genomes. GWAS analysis for PA, FA and PUC detected strong associations with two MYB genes similar to *MYB113* involved in the anthocyanin biosynthesis pathway and with a *COP1* gene, which encodes for a photo-regulatory protein and may be responsible for the PUC phenotype. Evidence was found of a duplication of an ancestral *MYB113* gene with a translocation from chromosome 10 to chromosome 1. Parental genotypes for the three genes were in agreement with the candidate genes identification performed in the S3MEGGIC population. Our new eggplant MAGIC population is the largest recombinant population in eggplant and is a powerful tool for eggplant genetics and breeding studies.

## Introduction

Multi-parent experimental populations are of great interest for the genetic dissection of quantitative traits as well as for the development of new recombinant materials for plant breeding (Huang et al., 2015). Despite their complex management and resources requirement, multi-parent advanced generation inter-cross (MAGIC) populations represent powerful next-generation mapping tools by combining high genetic diversity and recombination with low population structure (Arrones et al., 2020; Scott et al., 2020). MAGIC populations are already available in model species such as *Arabidopsis thaliana* and in several crops, such as cereals, pulses and vegetables (Kover et al., 2009; Bandillo et al., 2013; Pascual et al., 2015; Huynh et al., 2018), and have demonstrated their power to dissect the structure of complex traits (Dell’Acqua et al., 2015; Stadlmeier et al., 2018).

Although available MAGIC populations have become a useful resource for genetic studies and breeding, most of them have only exploited intraspecific variation. The incorporation of crop wild relatives (CWRs) as founders could be a way of including multiple wild genomic fragments or introgressions into cultivated background genomes (Arrones et al., 2020). Apart from being of great interest for genetic analysis, interspecific MAGIC populations can be useful for broadening the genetic base of crops and provide new variation for breeding multiple traits, including those related to adaption to climate change (Gramazio et al., 2020a). However, so far, the potential of interspecific MAGIC populations for plant breeding has largely remained unexploited (Arrones et al., 2020).

Eggplant (*Solanum melongena* L.) is a major vegetable crop of increasing importance, ranking fifth in global production among vegetables (Faostat, 2019). Despite its economic importance, eggplant has lagged behind other major crops and little efforts have been made to develop immortal experimental populations and genetic and genomic tools (Gramazio et al., 2018, 2019). So far, only one population of recombinant inbred line (RILs) and one set of introgression lines (ILs) are publicly available (Lebeau et al., 2013; Gramazio et al., 2017a), while no multiparent population has been developed so far. Conversely, in other related Solanaceae crops such as tomato, several experimental populations have been developed including MAGIC populations, which have allowed great advances in the genetic dissection of traits of interest (Pascual et al., 2015; Campanelli et al., 2019). For this reason, the development of this type of population would represent a landmark in eggplant breeding.

Anthocyanins represent a specific and relevant trait in eggplant which can thus be used as a model plant for other crops (Moglia et al., 2020). Anthocyanins play a key role in plant defence mechanisms and their synthesis and accumulation could vary in response to specific biotic and abiotic stresses (D’Amelia et al., 2018; Lv et al., 2019; Zhou et al., 2020). In addition, anthocyanins may prevent and alleviate human chronic diseases and provide health benefits (Toppino et al., 2020). In eggplant, anthocyanins are responsible for the purple colour of skin, one of the traits of greatest interest for eggplant breeding (Daunay and Hazra, 2012). Purple coloured eggplant fruits are the most demanded in many markets (Li et al., 2018), and developing dark purple-coloured eggplants, which result from the combination of anthocyanins with chlorophylls, is a major objective in eggplant breeding programmes. Eggplants are variable for the presence of anthocyanins not only in fruits, but also in other plant organs and tissues such as hypocotyl, stem, leaves, leaf veins, prickles, flower calyxes or corolla (Toppino et al., 2020). Anthocyanin biosynthesis has been widely studied in Solanaceae species (Van Eck et al., 1994; Borovsky et al., 2004; Gonzali et al., 2009), but its genetic control in eggplant has not been fully clarified. The anthocyanin biosynthetic pathway is a very conserved network in many plant species, where many enzymes and regulatory transcription factors (TFs) are involved (Albert et al., 2014).

The major anthocyanins in the eggplant fruit epidermis are delphinidin-3-p-coumaroylrutinoside-5-glucoside and the delphinidin-3-rutinoside (Mannella et al., 2012; Moglia et al., 2020). Fruit anthocyanins are highly dependent on light (Jiang et al., 2016), however, the genotypes that carry the PUC (pigmentation under the calyx) mutation are able to synthetize anthocyanins in the fruit epidermis independently of the incidence of light (Tigchelaar, et al., 1968). QTL-related studies using family-based or GWA mapping approaches evidenced that chromosome 10 harbours most of the QTL/genes involved in anthocyanin formation, distribution and accumulation (Doganlar et al., 2002; Barchi et al., 2012; Frary et al., 2014; Cericola et al., 2014; Toppino et al., 2016, 2020; Wei et al., 2020). The availability of high-quality eggplant genome sequences and transcriptomic data allowed the identification of putative candidate genes belonging to the MYB family controlling variation in anthocyanin content and fruit colour in eggplant (Docimo et al., 2016; Li et al., 2017, 2018; Xiao et al., 2018; Li et al., 2021; Moglia et al., 2020; Shi et al., 2021), highlighting their synteny with other Solanaceae. However, genes underlying the PUC phenotype have not been identified so far.

Here we report on the first eggplant MAGIC population derived from an interspecific cross of seven accessions of *S. melongena* and one of the wild relative *S. incanum* (Gramazio et al., 2019). It represents the largest experimental population described so far in eggplant, with a similar population size to MAGIC populations in other solanaceous crops. The population has been genotyped by applying the Single Primer Enrichment Technology (SPET) to explore its genetic architecture and the contribution of founders to the final population, and phenotyped for the presence of anthocyanins in the fruit epidermis and other plant organs and for the PUC phenotype. These traits were chosen due to their physiological, agronomic and morphological relevance, their high stability and heritability. An association analysis has been performed to locate genomic regions and to identify candidate genes involved in the traits under study.

## Results

### MAGIC population construction

Seven accessions of eggplant and one of the wild relative *S. incanum* were selected as founder parents (A-H) for the construction of the eggplant MAGIC population (S3MEGGIC). Following a funnel breeding scheme (Figure 1a and Figure S1) a total of 420 individuals of the MAGIC populations were obtained. First, founders were pairwise inter-crossed to produce four two-way hybrids (AB, CD, EF, and GH), which were subsequently inter-crossed in pairs to obtain four-way hybrids (ABCD and EFGH). One-hundred and forty-nine individuals of each of the two four-way hybrids were inter-crossed using a chain pollination scheme (Figure S1). Out of the theoretical maximum of 298 eight-way hybrid progenies (S0), seeds were obtained for 209 of them, of which 116 carried the *S. melongena* ASI-S-1 cytoplasm and 93 the *S. incanum* MM577 cytoplasm. Two plants per S0 progeny were used to advance the population reaching 402 S1 progenies. These S1 progenies were advanced through single seed descend (SSD) to obtain 391 S2 and 305 S3 MAGIC progenies. The final S3MEGGIC population was constituted by 420 S3 individuals, of which 348 individuals carried the cultivated cytoplasm and 72 the wild cytoplasm.

**Figure 1.**
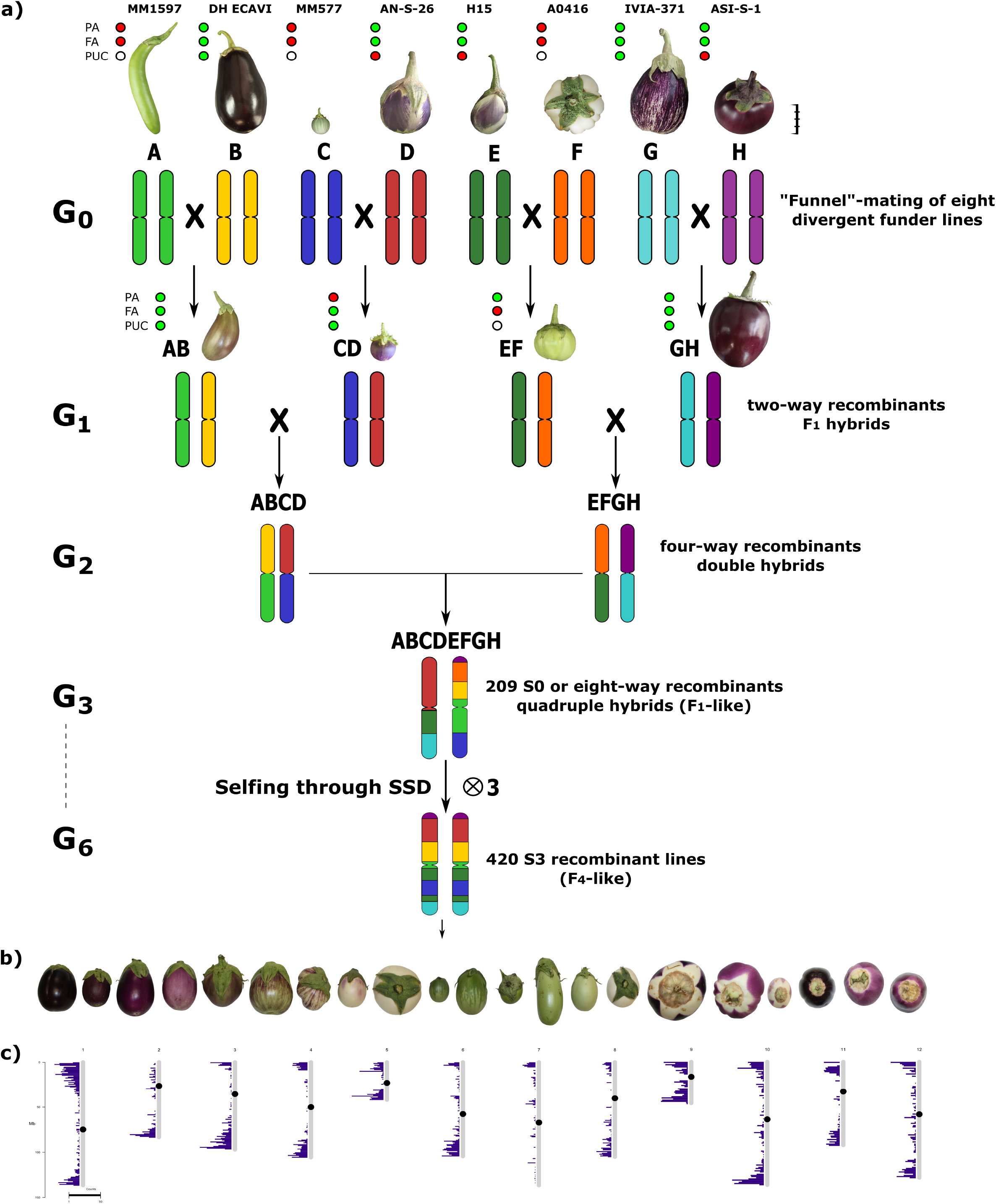
(a) The funnel breeding design used, across the six generations (G1 to G6), to develop the 420 S3 individuals of the S3MEGGIC population. The eight parents, coded from A to H and each with a different colour to represent their genomic background, are represented above at a scale based on the real fruit size. Scale bar represents 5 cm. The four two-way hybrids obtained in the G1 generation (AB, CD, EF and GH) are also represented at the same scale as the founders. Phenotyping of founders and two-way hybrids for absence (red) or presence (green) of plant anthocyanins (PA), fruit anthocyanins (FA) or pigmentation under the calyx (PUC). White dots for PUC mean uncertainty for non-anthocyanin fruits. (b) A representation of the phenotypic diversity of the S3 individuals found during the phenotyping. (c) The distribution of the molecular markers across the chromosomes used for the genotyping.

### SPET genotyping

The genotyping of the 420 S3 MAGIC individuals, the eight founders and the four two-way hybrids by the eggplant Single Primer Enrichment Technology (SPET) platform yielded 22,146 SNPs. After filtering, 7,724 high-confidence SNPs were retained for the subsequent analysis and the low percentage of missing calls (0.53%) was imputed. Filtered SNPs were distributed across the entire eggplant genome, although the distribution of SNPs varied within and among chromosomes (Table 1, Figure 1c). Chromosome 9 had the highest average marker density after SNP filtering with 173.07 SNPs per Mb, while chromosome 7 the lowest with an average of 23.03 SNPs per Mb. Generally, most of the SNPs were located in regions with high gene density and decayed around the centromere (Figure 1c). S3 MAGIC individuals exhibited a heterozygosity average of 6.87%, with only 15 individuals (3.57%) with a proportion of residual heterozygosity higher than 20% (Figure 2).

**Table 1.** Statistics of the genotyping using the eggplant SPET platform of the 420 S3MEGGIC population individuals using the ‘67/3’ eggplant reference genome (Barchi et al., 2019b).

| Chr | Markers | Filtered markers | % Markers | % Filtered markers | Chr length (Mb) | Marker density (Mb) | Filtered marker density |
| --- | --- | --- | --- | --- | --- | --- | --- |
| 1 | 3,234 | 1,205 | 14.60 | 15.60 | 13.66 | 236.82 | 88.24 |
| 2 | 1,119 | 424 | 5.05 | 5.49 | 8.34 | 134.21 | 50.85 |
| 3 | 2,004 | 758 | 9.05 | 9.81 | 9.71 | 206.41 | 78.07 |
| 4 | 1,567 | 588 | 7.08 | 7.61 | 10.57 | 148.23 | 55.62 |
| 5 | 1,438 | 571 | 6.49 | 7.39 | 4.39 | 327.80 | 130.16 |
| 6 | 2,108 | 794 | 9.52 | 10.28 | 10.90 | 193.38 | 72.84 |
| 7 | 2,483 | 328 | 11.21 | 4.25 | 14.24 | 174.34 | 23.03 |
| 8 | 1,392 | 550 | 6.29 | 7.12 | 10.96 | 126.98 | 50.17 |
| 9 | 1,841 | 625 | 8.31 | 8.09 | 3.61 | 509.80 | 173.07 |
| 10 | 2,069 | 798 | 9.34 | 10.33 | 10.67 | 193.86 | 74.77 |
| 11 | 1,260 | 453 | 5.69 | 5.86 | 7.23 | 174.23 | 62.64 |
| 12 | 1,631 | 630 | 7.36 | 8.16 | 10.05 | 162.33 | 62.70 |
| Total | 22,146 | 7,724 | 100.00 | 100.00 | 114.33 |  |  |
| Average |  |  |  |  |  | 215.70 | 76.85 |

**Figure 2.**
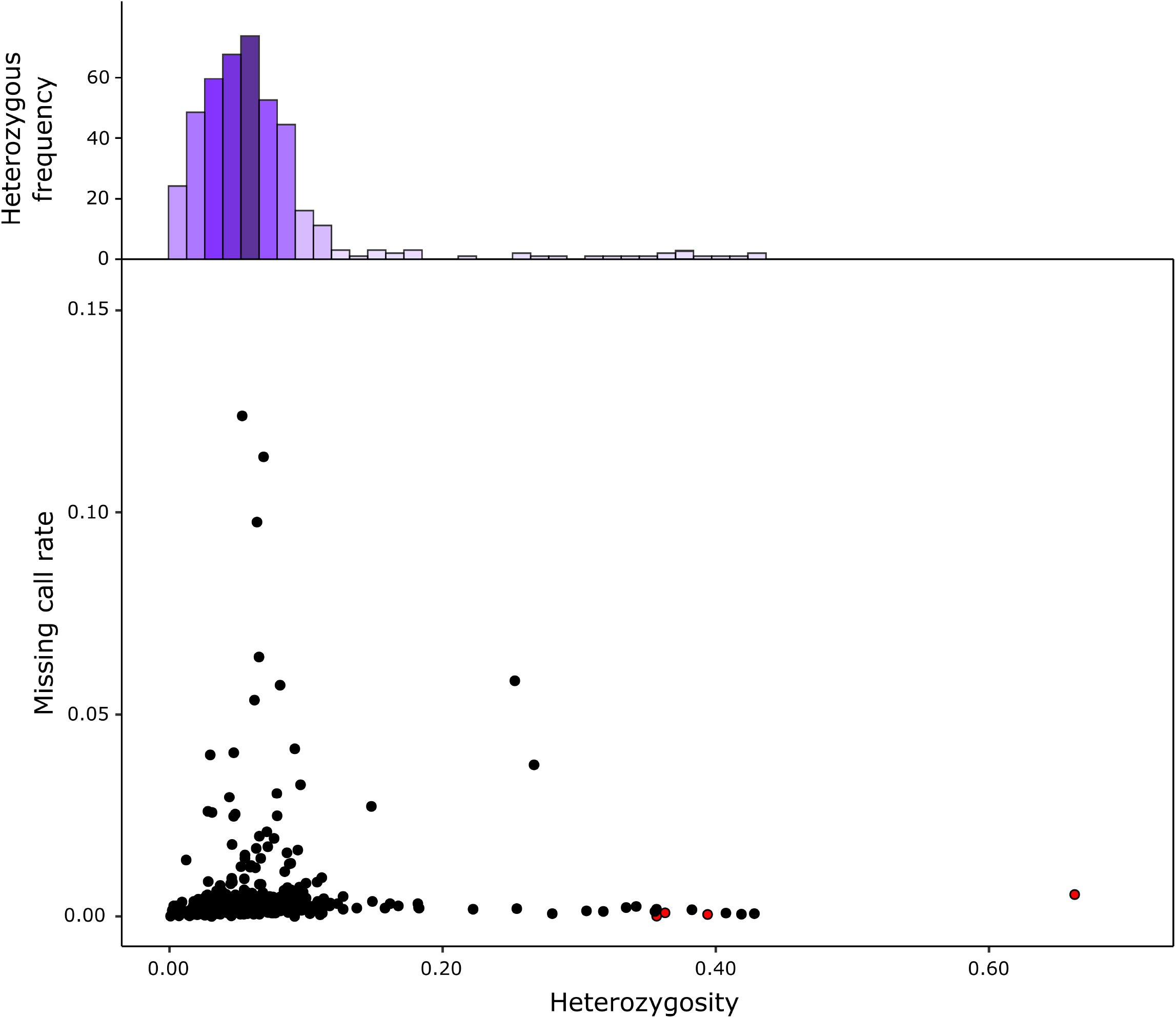
Heterozygosity and proportion of missing data from the S3MEGGIC population. The lower graph reports residual heterozygosity of S3 individuals as black dots compared against two-way hybrids represented as red dots. The top histogram represents the observed heterozygosity which is skewed to the left (mean 6.87%; mode 5.02%).

### Population structure

Population stratification performed by PCA indicated the absence of subgroups in the S3 individuals since no clear clustering was observed (Figure 3). The first two principal components (PCs) explained respectively 5.15% (PC1) and 3.30% (PC2) and the first 10 PCs explain altogether only 25.14% of the total variation, revealing the absence of genetic structure in the S3MEGGIC population. No differentiated clusters among individuals carrying the wild (*S. incanum* MM577) or the cultivated (*S. melongena* ASI-S-1) cytoplasm were observed. An Analysis of Molecular Variance (AMOVA) was also performed revealing that only 0.29% of the total sums of squares is accounted for the molecular variation among the *S. melongena* and *S. incanum* cytoplasm groups resulting in a very small phi-value of 0.0019, which indicates a low level of differentiation supporting that no population structure exists.

**Figure 3.**
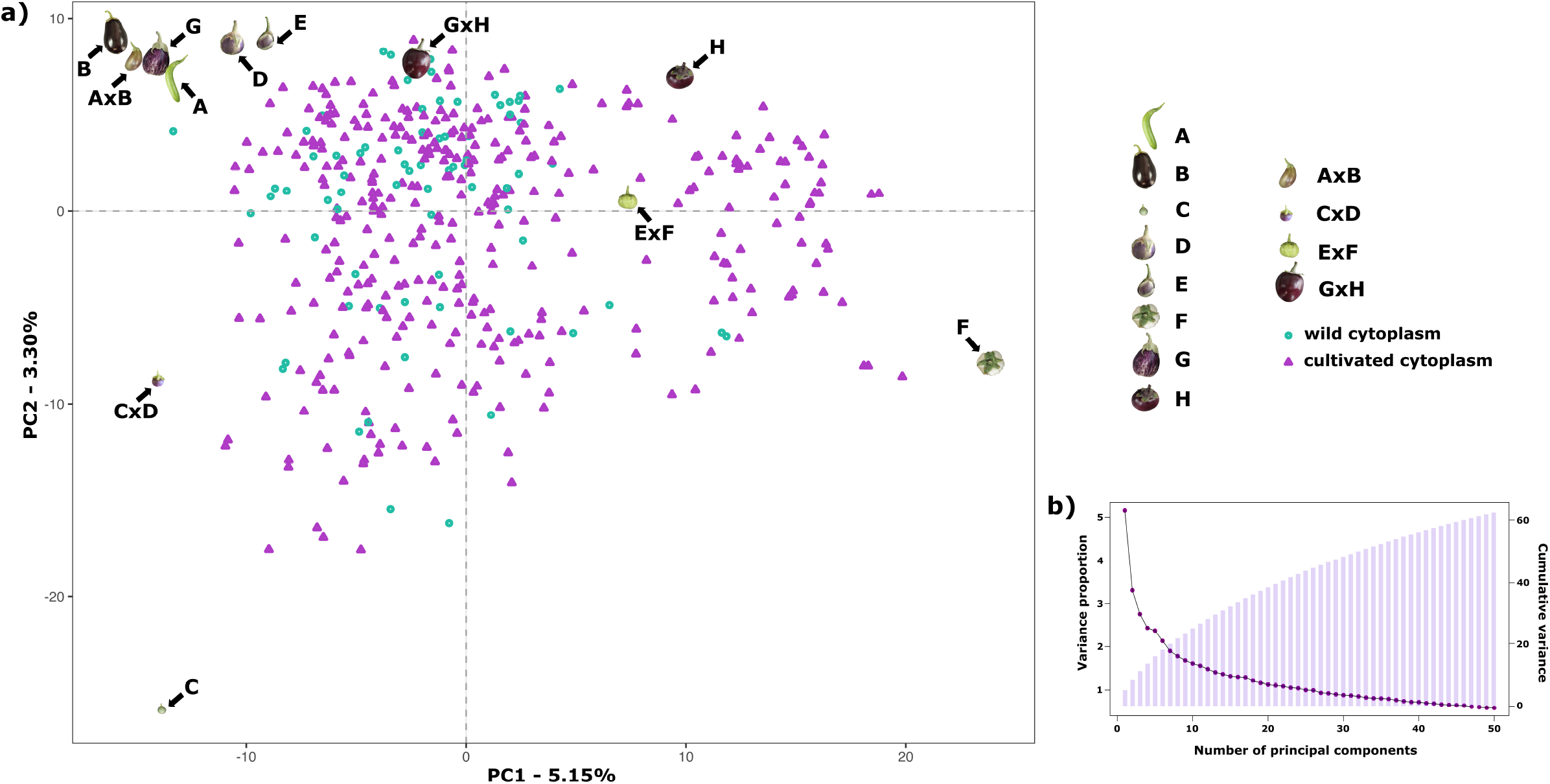
Results of a Principal Component Analysis (PCA) on the S3MEGGIC population. (a) PCA of the first two PCs including S3 individuals, founder lines and two-way hybrids. S3 individuals with wild cytoplasm are represented with blue dots while those ones with cultivated cytoplasm are represented with purple triangles. (b) Scree plot of the PCs (x-axis) and their contribution to variance (left y-axis); bar blot of the PCs (x-axis) and the cumulative proportion of variance explained (right y-axis).

The genome mosaics reconstruction of the S3 MAGIC individuals in terms of the eight founder haplotypes showed different haplotype block proportions depending on the genomic position for all chromosomes (Figure 4). The estimated contribution of some founders to the overall S3MEGGIC population differed from the expected value of 12.5%. Two of the founder genomes (A0416 and IVIA-371) had a high representation in the genome of the S3 individuals (32.6% and 23.6%, respectively) while two others (AN-S-26 and H15) had a small representation (0.3% in both cases). The wild founder *S. incanum* had an average haplotype representation of 5.8%.

**Figure 4.**
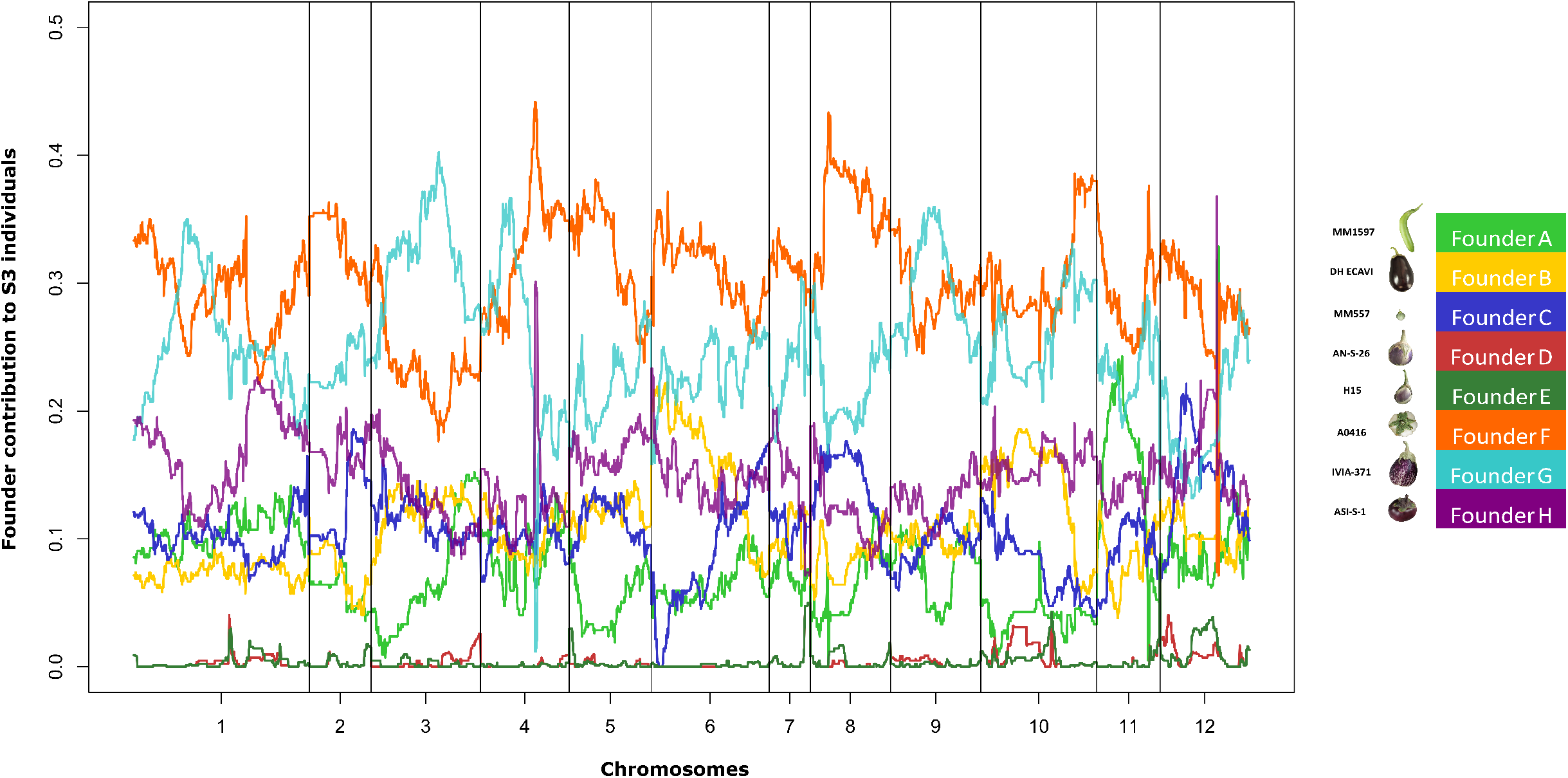
Genome-wide founder haplotype blocks assignment across the entire S3MEGGIC population. In the x-axis is represented the 12 eggplant chromosomes and in the y-axis the average percentage of founders contribution for the 420 S3 individuals. In the legend the color code associated with each founder as in Figure 1.

### Phenotypic variation among MAGIC individuals and association analysis

The screening for absence or presence of plant (PA) and fruit (FA) anthocyanins and pigmentation under the calyx (PUC) in purple fruits of the 420 S3 MAGIC individuals revealed considerable variation (Figure 1b). Out of the 420 S3 individuals, 57.6% displayed PA and 37.5% FA. Among individuals displaying FA, 64.3% showed PUC.

Given that no population structure was observed for the S3MEGGIC population, the phenotypic data together with the genotypic information was used for Genome-Wide Association Study (GWAS) analysis (Figure 5). GWAS was performed taking into account the kinship in a mixed linear model (MLM) leading to the identification of significant associations for the evaluated traits.

**Figure 5.**
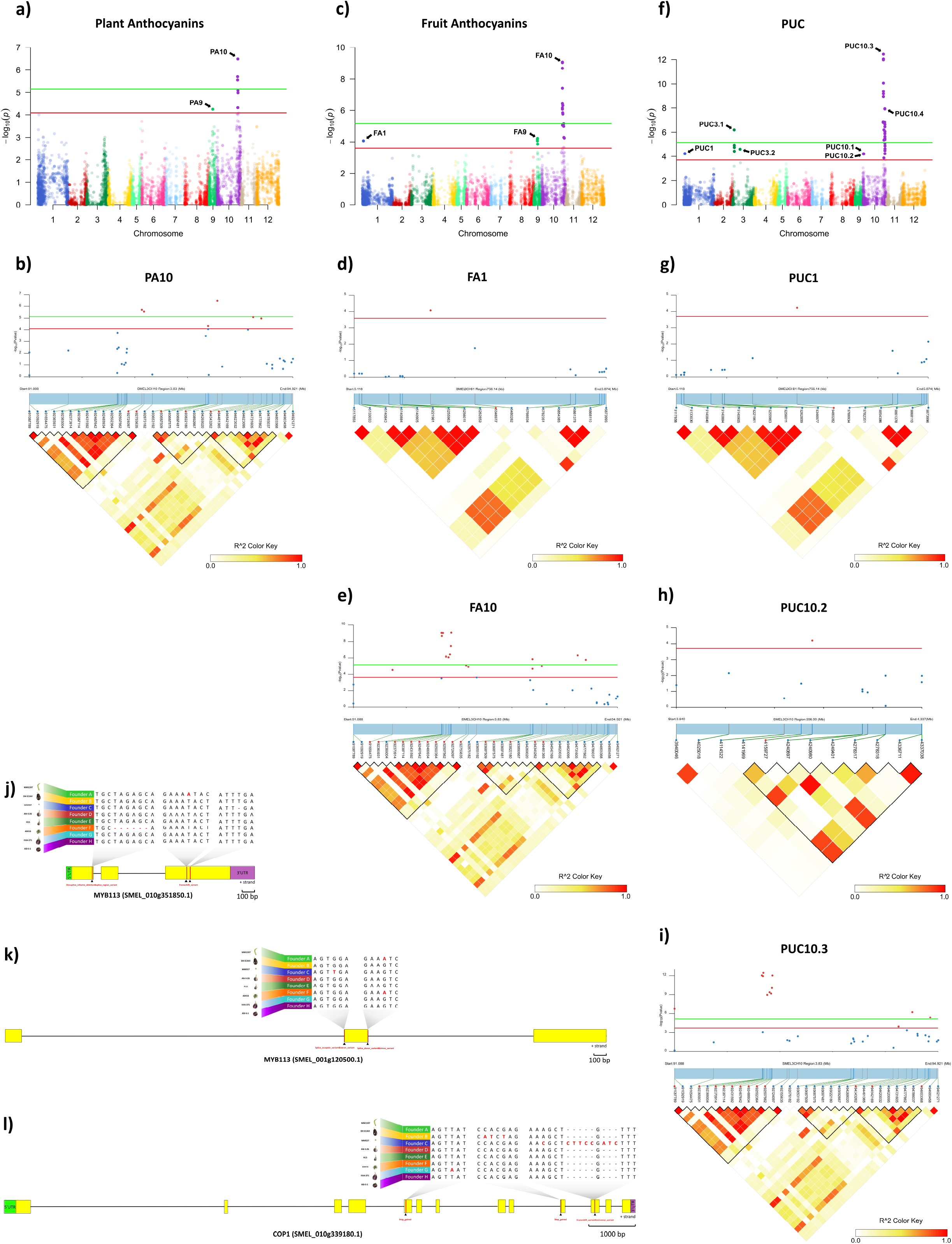
Genome-wide association mapping with LD block heatmap of significant regions and potential candidate genes controlling plant anthocyanins (PA), fruit anthocyanins (FA) and anthocyanin pigmentation under the calyx (PUC) in eggplant. (a, c, f) Manhattan plot for PA (a), FA (c) and PUC (f). Arrows indicate the position of main peaks detected for each trait. The red and green horizontal lines represent, respectively, FDR and Bonferroni significance thresholds at p=0.05. (b, d, e, g, h, i) Local Manhattan plot (top) and LD heatmap (bottom) surrounding the peaks PA10 (b), FA1 (d), FA10 (e), PUC1 (g), PUC10.2 (h) and PUC10.3 (i). Pairwise LD between SNPs is indicated as values of *R*^*2*^ values: red indicates a value of 1 and white indicates 0. (j-l) Structure of candidate genes MYB113 (SMEL_010g351850.1) (j), MYB113 (SMEL_001g120500.1) (k) and COP1 (SMEL_010g339180.1) (l) and effect of the high-impact variants detected in those of the eight founders.

#### Plant Anthocyanins (PA)

The Manhattan plot for PA revealed two major peaks on chromosome 9 and 10 (PA9 and PA10), with seven significant SNPs over the FDR threshold (LOD > 4.09) and three of them over the Bonferroni threshold (LOD > 5.16) (Figure 5a). One SNP (LOD = 4.25) mapped on PA9 region (between 17.0 and 17.4 Mb; Figure S2a), and six SNPs, with a LOD between 4.31 and 6.48, were mapped on PA10 region (between 91.08 and 94.81 Mb; Figure 5b).

#### Fruit Anthocyanins (FA)

For FA, 22 significant SNPs above FDR threshold (LOD > 3.6), which included 11 SNPs above the Bonferroni threshold (LOD > 5.16), were plotted on three major peaks located on chromosomes 1, 9 and 10 (FA1, FA9 and FA10, respectively) (Figure 5c). One SNP (LOD = 4.07) was detected on FA1 region (between 5.11 and 5.88 Mb) in position 5,346,977 (Figure 5d). Five SNPs, with a LOD between 3.86 and 4.22, were located on FA9 region (between 16.2 and 17.4 Mb; Figure S2b), which overlapped with the PA9 region found for PA (Figure S2a). The other sixteen SNPs (LOD between 4.52 and 9.06) were detected on FA10 region, between 91.08 and 94.81 Mb (Figure 5e), corresponding to the PA10 region where significant associations were detected for PA (Figure 5b).

#### Pigmentation Under the Calyx (PUC)

Among the three traits evaluated, PUC was the one with the highest number of significant associations. The Manhattan plot for PUC revealed seven main peaks on chromosomes 1 (PUC1), 3 (PUC3.1 and PUC3.2) and 10 (PUC10.1-10.4), with 36 SNPs over the FDR threshold (LOD > 3.7) and 23 of them above the Bonferroni threshold (LOD > 5.16) (Figure 5f). One SNP with a LOD of 4.21 was located on the PUC1 region (between 5.11 and 5.88 Mb) in position 5,480,282 (Figure 5g), 133,305 bp away of the significant SNP on the FA1 region (Figure 5d). On chromosome 3, four SNPs (LOD between 4.41 and 6.18) were detected in PUC3.1 region (between 7.22 and 8.65 Mb; Figure S2c), and one SNP (LOD = 4.58) in PUC3.2 region (between 31.54 and 40.09 Mb; Figure S2d). Thirty SNPs were mapped on chromosome 10 in four genomic regions. One of the SNPs (LOD = 4.2) was identified in PUC10.1 region (between 2.08 and 2.67 Mb) in position 2,388,375 (Figure S2e), and other one (LOD = 4.2) in PUC10.2 region (between 3.94 and 4.34 Mb; Figure 5h). Other twelve SNPs were located in PUC10.3 region, between 91.08 and 94.81 Mb (LOD between 3.86 and 12.44; Figure 5i), as observed for PA (Figure 5b) and FA (Figure 5d). The rest of 16 SNPs (LOD between 3.71 and 7.93) were found in PUC10.4 region, between 98.25 and 100.55 Mb (Figure S2f).

### Candidate genes for anthocyanin biosynthesis

Based on the results of the GWAS analysis, putative candidate genes were identified close to or within LD blocks defined in the genomic regions with significant associations (Table S1). On chromosome 10, in the genomic region (91.08 - 94.81 Mb) associated with all the evaluated traits (PA10, FA10 and PUC10.3), a candidate gene was identified as similar to *MYB113* (SMEL_010g351850.1), a well-known regulatory transcription factor controlling anthocyanin synthesis in eggplant (Zhou et al., 2019; Shi et al., 2021). In addition, in the genomic region associated with FA and PUC (FA1 and PUC1) on chromosome 1 (5.11 - 5.88 Mb), another candidate gene was identified similar to *MYB113* (SMEL_001g120500.1). Variants that predicted high impact effects on protein function were annotated by SnpEff for both *MYB113* genes in the population founders (A, C and F) that do not present anthocyanins in plants and fruits, as confirmed by aligning the founder gene sequences (Figure 5j and 5k). Specifically, for the founders A and C, single frameshift variants were identified in two different positions while a disruptive inframe deletion and a slice region variant were identified for the founder F in a third region of the eggplant gene SMEL_010g351850.1 (Figure 5j). For SMEL_001g120500.1, founders A and F exhibited the exact same splice donor and intron variant predicted as high impact, while in the founder C a splice acceptor and intron variant were identified (Figure 5k). Founder reconstruction of the S3 individuals and haplotype blocks were estimated for both candidate gene regions (chromosome 10: 91.08 - 94.81 Mb; chromosome 1: 5.11 - 5.88 Mb) (Figure S3a and S3b). Founders that most contribute to anthocyanin-related traits in these regions are IVIA-371 (111 and 97 S3 individuals, respectively) and ASI-S-1 (74 and 75 S3 individuals, respectively). Furthermore, a reciprocal best hit BLAST analysis of the SMEL_001g120500.1 onto the tomato genome indicated that this gene corresponded to the orthologue *SlANT1*-like with 70.99% identity. The eggplant SMEL_010g351850.1 corresponded to the tomato orthologue *SlAN2*-like with a 73.73% identity, which has been described as the best candidate gene for anthocyanin fruit biosynthesis in tomato (Yan et al., 2020). These results suggest that a single duplication event and a translocation of a fragment of at least 357 kb from chromosome 10 to 1 occurred during eggplant evolution (Figure 6).

**Figure 6.**
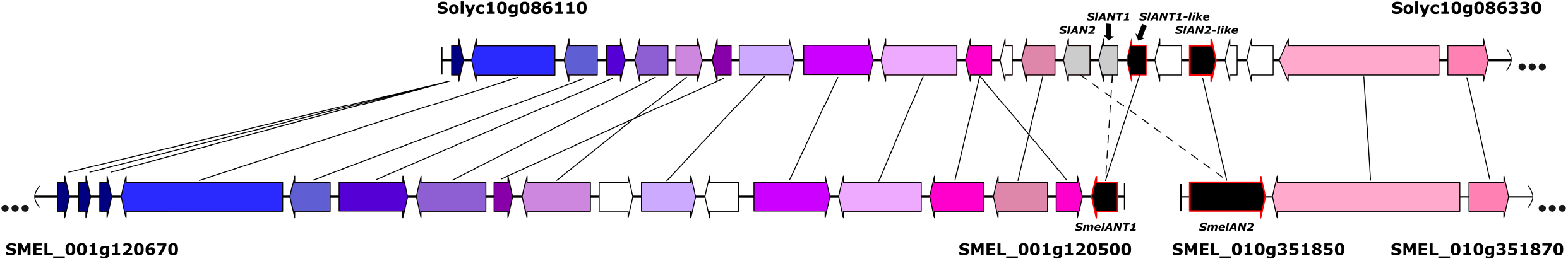
Tomato-eggplant microsynteny representation of a tomato region from chromosome 10 (Solyc10), eggplant region of chromosome 10 (SMEL_010) and a 357 kb fragment of eggplant chromosome 1 (SMEL_001), where candidate genes for anthocyanins synthesis are located.

A candidate gene that corresponds to a *COP1* (SMEL_010g339180.1), a gene that encodes for photo-regulatory proteins (Jiang et al., 2016; Li et al., 2018; He et al., 2019; Naeem et al. 2019), was reported in the genomic regions associated with PUC (PUC10.2) on chromosome 10 (3.94 - 4.34 Mb; Table S1). High-effect variants were detected in *COP1* gene in those founders able to synthesize anthocyanin under the calyx (B and G) but also for the green wild species *S. incanum* (C) (Figure 5l). Considering PUC as dominant, the green C founder mutation was confirmed by the F1 hybrid phenotype from the C x D inter-cross, *PUC* (pigmentation under the calyx) x *puc* (no anthocyanins under calyx), which showed anthocyanin fruits with pigmentation under the calyx (Figure 1a). The wild founder presented multiple SNPs compared with the rest that were predicted to cause a frameshift and missed variants, while founders B and G exhibited variants that were predicted to produce stop codons (Figure 5l). Founders with higher haplotype representation contributing to PUC in this region (chromosome 10: 3.94 - 4.34 Mb) are IVIA-371 (113 S3 individuals) and DH_ECAVI (71 S3 individuals) (Figure S3c).

Furthermore, close to LD blocks in the PA9 and FA9 (between 17,862,090 and 17,872,428 Mb) and PUC10.1 (between 3,128,678 and 3,143,068 Mb) regions, we found two candidate genes annotated as similar to *BHLH*, basic helix loop helix protein A (SMEL_009g326640.1), and similar to *SPA3*, protein SPA1-RELATED 3 (SMEL_010g338090.1), respectively (data not shown). Although they were described as genes related to the anthocyanin biosynthetic pathway (Maier et al., 2013; Li et al., 2018; Shi et al., 2021), no high-effects variants were identified in these genes for any of the founders.

## Discussion

MAGIC populations are outstanding genetic materials for identifying gene-trait associations with high resolution (Arrones et al., 2020; Scott et al., 2020). The introduction of multiple founders with increased genetic and phenotypic diversity together with the multiple rounds of inter-crossing and selfing increases the number of accumulated recombinant events and, thus, improving mapping accuracy (Scott et al., 2020). By introducing a wild relative as a founder parent, the genetic variability in the population increases, which is a key point for QTL identification (Gramazio et al., 2020a). Here we present the first eggplant MAGIC population of which one of the founder was one accession of the wild close relative *S. incanum*.

Large population sizes are essential to increase power and mapping resolution in MAGIC populations (Collard et al., 2005; Valdar et al., 2005; Jaganathan et al., 2020). Following a simple “funnel” scheme design, the population was kept as large as possible to gather a large number of recombination events. However, a sharp reduction in the number of progenies was observed at the S0 generation, which might be related to the use as a founder of the wild species *S. incanum* (C) as a female parent to obtain the simple (CD) and the double (ABCD) hybrids. This interspecific crossing dragged the maternal cytoplasmic background of the wild parent, which might have caused partial sterility and bias in subsequent generations. Some studies confirmed a strong effect of wild *Solanum* cytoplasm in the reduction in pollen fertility of the alloplasmic lines (Isshiki et al., 2020, Khan et al., 2020). However, the PCA highlighted the absence of population structure, also confirmed by the lack of genetically differentiated cytoplasmic groups.

The genotyping of the S3MEGGIC population was carried out with the 5k probes eggplant SPET platform with a well-distributed marker density along all chromosomes (Barchi et al., 2019a). This genotyping strategy has already been used in the analysis of biparental populations (Herrero et al., 2020), and here we have verified that its use can be extended to multiparent populations. The genotyping revealed a low heterozygosity in the S3MEGGIC individuals, similar to the expected value for a F5-like biparental inter-cross generation (6.25%). The contributions of each of the founder parents to the S3MEGGIC population revealed that some parents had higher representation than others. Apart from drift effects, several biological reasons could potentially explain cryptic selection processes causing the unbalanced representation of the genomes (Rockman and Kruglyak, 2008; Thépot et al., 2015). Among them, seed dormancy, delayed germination, precocity, reduced fertility, parthenocarpy associated to some genomes, which have already been reported in eggplant and other crops (Barchi et al., 2010; Khan et al., 2015; Prohens et al., 2017). The rather limited contribution of the wild species *S. incanum* to the final S3 MAGIC individuals may have been caused by selection pressure, as progenies bred from crosses involving two different species tend to suffer from reduced fertility and show segregation distortion (Lefebvre et al., 2002; Barchi et al., 2010). In addition, *S. incanum* has a recalcitrant germination and a very erratic flowering and fruit set, which strongly depends on environmental conditions (Gisbert et al., 2011; Mangino et al., 2020, 2021). Other reasons for the segregation distortion could be the inability of the current genotyping density to efficiently distinguish between founders that are genetically closer like AN-S-26 and H15 genotypes. This phenomenon has already been observed in previous MAGIC populations (Dell’Acqua et al., 2015). A deeper genotyping resulting in a better haplotype reconstruction might shed light on the mechanisms that have led to the unbalanced representation of the founder parents genome in the S3MEGGIC population.

The anthocyanin biosynthetic pathway is one of the most studied biochemical routes in plants, given its physiological importance. Major structural genes of this pathway are under the control of a regulatory complex, where myeloblastosis (MYB) TFs are recognized as main regulators, alone or in complexes with other TFs (Ramsay and Glover, 2005; Kiferle et al., 2015; Liu et al., 2018). Activator or repressor MYB proteins directly, and competitively, bind the basic-helix-loop-helix (bHLH) via the amino terminus domain, and can act as positive or negative transcriptional regulators in a tissue-specific mode to modulate anthocyanin synthesis (Barchi et al., 2019b; Moglia et al., 2020). In eggplant, anthocyanins related MYBs protein-encoding genes have been reported as related with fruit peel coloration (Zhang et al., 2014; Docimo et al., 2016; Xiao et al., 2018; Moglia et al., 2020; Toppino et al., 2020). In tomato, a cluster of four different MYB proteins was reported as involved in the anthocyanin synthesis located on chromosome 10 and encoded by *SlAN2, SlANT1, SlANT1*-like and *SlAN2*-like genes (Solyc10g086250, Solyc10g086260, Solyc10g086270, Solyc10g086290, respectively); however genetic associations to only two paralogue *MYB113* genes were detected in the S3MEGGIC population on chromosomes 1 and 10 (SMEL_001g120500.1 and SMEL_010g351850.1). The same occurs in potato, in which only two anthocyanin genes have been identified, i.e. Sotub10g028550 and Sotub10g028540, both located on chromosome 10. While in pepper, in addition to CA10g11690 and CA10g11650, a third gene (CA10g11710) has been identified as an orthologue to *SlAN2*-like (Barchi et al., 2019b). In eggplant, these two tomato orthologs were previously described as *SmelANT1* and *SmelAN2* (Docimo et al., 2016; Barchi et al., 2019b), corresponding to *SlANT1* and *SlAN2*. However, the reciprocal best hit BLAST analysis of the eggplant coding proteins showed stronger homology respectively to tomato *SlANT1*-like and *SlAN2*-like. Although previous studies in eggplant indicated that overexpression of *SmelANT1* accounts for constitutive upregulation of most anthocyanin biosynthetic genes (Zhang et al., 2014; Shi et al., 2021), we considered *SmelAN2* as the best candidate gene for different reasons. Previous studies in eggplant located major anthocyanin-related QTLs on chromosome 10 (Barchi et al., 2012; Cericola et al., 2014; Toppino et al., 2016), which is in agreement with the highest association signals for anthocyanin related traits in our GWAS results. Furthermore, recent studies in tomato suggested that *SlAN2*-like functions as an activator to regulate biosynthesis genes, including *SlANT1*-like, and controls the accumulation of anthocyanins (Yan et al., 2020). However, in the S3MEGGIC population, high-impact variants on protein function were found in both *MYB113* genes for non-anthocyanin fruits. These results could suggest that a duplication of function occurred during eggplant evolution and both genes may be required for anthocyanin synthesis.

Although it has been demonstrated that the activation of the anthocyanin biosynthetic pathway in eggplant is strongly regulated by light (Li et al., 2018; Xiao et al., 2018), the PUC mutation confers to some genotypes the ability to synthesize anthocyanins under the calyx regardless of light. PUC has been described as a single gene independent of the presence or absence of anthocyanins in the fruit (Tigchelaar, et al., 1968). In this study, a candidate gene related with light-dependent anthocyanin biosynthesis in fruits was detected at the beginning of chromosome 10. The constitutive photomorphogenic1 (*COP1*) gene has been reported as a regulatory TF responsible for mediating light-regulated gene expression and development (Jiang et al., 2016; Li et al., 2018; He et al., 2019; Naeem et al. 2019). *COP1* has been demonstrated to act as a light-inactivable repressor interacting with MYB TFs (Jiang et al., 2016), and it is considered a ‘molecular switch’ in metabolic processes which are stimulated by light. In presence of light, *COP1* expression is inhibited, and its concentration decreased rapidly promoting anthocyanins synthesis; while under dark conditions, *COP1* expression is induced and promotes the degradation of the photomorphogenesis-promoting TFs and MYBs inhibition by conforming the *COP1*/Suppressor of phya□105 (*SPA*) ubiquitin ligase complex (Maier et al., 2013; Jiang et al., 2016, Li et al., 2018). Although PUC can only be observed in anthocyanic fruits, high-impact variants were also identified in the wild species *S. incanum COP1* gene (founder C, green fruit). These results suggest that the dominant allele *PUC* is probably the ancestral gene and *puc* is a derived loss-of-function allele that appeared during the eggplant domestication and crop evolution. The ancestral allele is more common in eggplants from western countries, where European markets demand for more homogeneously pigmentation eggplants. On the contrary, in Asian eggplants it is more common to find the *puc* phenotype, such as in the ASI-S-1 founder.

In conclusion, the S3MEGGIC population represents a landmark breeding material and tool of great value which allows the study and fine mapping of complex traits due to: i) the highly phenotypically diverse founders; ii) the large population size being the largest developed eggplant experimental population up to now; iii) the high degree of homozygosity of the final individuals, which constitute a population of fixed “immortal” lines nearly homozygous at each locus; and, iv) the tailored genotyping SPET platform used for the genetic analysis of the population, which has been developed from the WGS of the founders and allows the comparison with materials genotyped with the same setup (Gramazio et al., 2020b). In addition, the S3MEGGIC populaton has demonstrated its potential usefulness for association studies allowing the establishment of marker associations to anthocyanin-related genes and identification of candidate genes for plant and fruit anthocyanins, including the first identification of a candidate gene for an economically relevant breeding trait in eggplant such as PUC.

## Experimental procedures

### MAGIC population construction

The eggplant MAGIC population has been developed by intermating seven cultivated eggplants, i.e. MM1597 (A), DH ECAVI (B), AN-S-26 (D), H15 (E), A0416 (F), IVIA-371 (G) and ASI-S-1 (H), and the *S. incanum* accession MM577 (C) (Figure 1a). The wild relative founder was chosen for its tolerance to some biotic and abiotic stresses, mainly drought (Knapp et al., 2013), and for showing high phenolic content (Prohens et al., 2013). The performance of the founders was comprehensively characterized in previous morphoagronomic and genetic diversity studies (Hurtado et al., 2014; Gramazio et al., 2017b; Kaushik et al., 2018) and their genomes have been resequenced (Gramazio et al., 2019). The latter study highlighting that in the founder parents the residual heterozygosity was less than 0.06%.

In order to develop the eggplant S3MEGGIC (S3 Magic EGGplant InCanum) population, founder lines have been inter-crossed by following a simple “funnel” approach (Wang et al., 2017; Arrones et al., 2020) (Figure 1a). The eight founders (A-H) have been pairwise inter-crossed to produce two-way or simple F1 hybrids (AB, CD, EF, GH), which were subsequently inter-crossed in pairs (AB × CD and EF × GH) to obtain two four-way or double hybrids (ABCD and EFGH). In order to achieve a complete admixture of all founder genomes and to avoid assortative mating, the double hybrids were intercrossed following a chain pollination scheme, with each individual being used as female as well as male parents (Díez et al., 2002) (Figure S1a).

All the eight-way or quadruple hybrids obtained (S0 generation) presented all the eight genomes randomly shuffled and only differed for the cytoplasm inherited from the maternal parent. The S0 progenies obtained using the double hybrid ABCD as female parent carried the cytoplasm of the wild *S. incanum* MM577, while those derived using the double hybrid EFGH as female parent carried the cytoplasm of the cultivated *S. melongena* ASI-S-1. Subsequently, the S0 progenies were selfed for three generations by single seed descent (SSD) to obtain the S3 segregating individuals that were phenotyped and genotyped in this study. To ensure the continuity of the S0 progenies and to accelerate the self-fertilization process, four plants of each S0 progeny were germinated, selecting for the next generation (S1) only the first two that set viable seed (Figure S1b). From each of the two S0 selected plants, two S1 plants were germinated and only the first one setting fruit was selected for the S2 generation. The same was done for the S3 generation, so that for each progeny two plants were germinated and phenotyped but only one was used for originating the next generation. Differently, when in the S3 progenies two individuals displayed some phenotypic differences, both were included in the S3MEGGIC population.

### Cultivation conditions

Seeds were germinated in Petri dishes, following the protocol developed by Ranil et al. (2015), and subsequently transferred to seedling trays in a climatic chamber under photoperiod and temperature regime of 16 h light (25 °C) and 8 h dark (18 °C). After acclimatization, plantlets were transplanted to 15 L pots and grown in a pollinator-free benched glasshouse of the UPV, Valencia, Spain (GPS coordinates: latitude, 39° 28′ 55″ N; longitude, 0° 20′ 11″ W; 7 m above sea level). Plants were spaced 1.2 m between rows and 1.0 m within the row, fertirrigated using a drip irrigation system and trained with vertical strings. Pruning was done manually to regulate vegetative growth and flowering. Phytosanitary treatments were performed when necessary. In order to shorten generation time of subsequent generations (S0-S3), plantlets were transplanted to individual thermoformed pots (1.3 L capacity) in a pollinator-free glasshouse and selfings were stimulated by mechanical vibration.

### High-throughput genotyping

Young leaf tissue was sampled from 420 S3 individuals, the eight founders and the four two-way hybrids. Genomic DNA was extracted using the SILEX extraction method (Vilanova et al., 2020) and checked for quality and integrity by agarose electrophoresis and Nanodrop ratios (260/280 and 260/230), while its concentration was estimated with Qubit 2.0 Fluorometer (Thermo Fisher Scientific, Waltham, MA, United States). After dilution, the samples were sent to IGA Technology Services (IGATech, Udine, Italy) for library preparation and sequencing with NextSeq500 sequencer (150 paired-end) for high-throughput genotyping using the Single Primer Enrichment Technology (SPET) technology using the 5k probes eggplant SPET platform (Barchi et al., 2019a). The latter comprises 5,093 probes, and was developed by filtering out the most informative and reliable polymorphisms (3,372 of them in CDSs and 1,721 in introns and UTRs regions) from the set of over 12 million SNPs identified among the MAGIC founders (Gramazio et al., 2019).

Raw reads were demultiplexed and the adapters removed using standard Illumina pipeline and Cutadapt (Martin, 2011) while trimming was performed by ERNE (Del Fabbro et al., 2013). Clean reads were mapped onto the eggplant reference genome “67/3” (Barchi et al. 2019b) using BWA-MEM (Li, 2013) with default parameters and only uniquely aligned reads were selected for the variant calling performed with GATK 4.0 (DePristo et al., 2011) following the best practice recommended by the Broad Institute (http://www.broadinstitute.org).

The SNPs identified by SPET were filtered using the Trait Analysis by aSSociation, Evolution and Linkage (TASSEL) software (ver. 5.0, Bradbury et al., 2007) in order to retain the most reliable ones (minor allele frequency > 0.01, missing data < 10% and maximum marker heterozygosity < 70%). In addition, a LD k-nearest neighbour genotype imputation method (LD KNNi) was performed to fill missing calls or genotyping gaps.

### Population structure, heterozygosity, haplotype blocks inferring

A principal component analysis (PCA) was performed to assess the population structure of S3MEGGIC using the R package vcfR (Knaus and Grünwald, 2017) and the function glPCA of the Adegenet package (Jombart, 2008). Finally, the PCA was graphically plotted with ggplot2 (Wickham, 2016). An Analysis of Molecular Variance (AMOVA) was performed to estimate population differentiation according to the cytoplasm (cultivated vs. wild) of the individuals of the S3MEGGIC population by using the function poppr.amova of the poppr R package (Kamvar et al., 2014). The residual heterozygosity and its distribution were evaluated with TASSEL software (ver. 5.0, Bradbury et al., 2007). Parental contribution to S3MEGGIC individuals and haplotype blocks were estimated by using R-package HaploBlocker (Pook et al., 2019).

### Phenotyping and Genome-Wide Association Study (GWAS)

Phenotypic data were collected from the 420 S3 individuals grown during the 2019/2020 season. Three traits were screened using a binary classification (presence/absence): anthocyanins in vegetative plant tissues (PA), anthocyanins in fruit epidermis (FA) and anthocyanin pigmentation under the calyx (PUC) (Figure S4). The Presence of PA was phenotyped when purple coloration was observed in any plant parts such as in stem, branches, leaf veins or prickles. For FA, anthocyanins were considered as present when purple coloration was observed in the fruit epidermis regardless of their distribution (uniform, listed, etc.) or intensity. PUC could only be phenotyped in anthocyanic fruits by removing the calyx and observing the presence of anthocyanins under it. Traits were screened at the stage of commercial maturity (e.g., when the fruit was physiologically immature) which is the best stage for phenotyping these traits. Phenotypic characteristics of the eight founders and two-way hybrids are described in Figure 1a.

Using the phenotypic and genotypic data collected of the S3MEGGIC individuals, a Genome-Wide Association Study (GWAS) was performed for the selected traits using the TASSEL software (ver. 5.0, Bradbury et al., 2007). For the association study, mixed linear model (MLM) analyses were conducted. MLM analysis uses both fixed and random effects which incorporates kinship among the individuals. The multiple testing was corrected with the Bonferroni and the false discovery rate (FDR) methods (Holm, 1979; Benjamini and Hochberg, 1995) to identify candidate associated regions at the significance level of 0.05 (Thissen et al., 2002). SNPs with LOD (-log10(p-value)) over these thresholds or cut off values were declared significantly associated with anthocyanin presence. The R qqman package (Turner, 2018) was used to visualize the Manhattan plots and LDBlockShow (Dong et al., 2020) to determine linkage disequilibrium (LD) and plot haplotype block structure. The pattern of pairwise LD between SNPs was measured by LD correlation coefficient (r^*2*^), considering haplotype blocks with default r^*2*^ values greater than 0.5 and supported by the solid spine of LD method (Gabriel et al., 2002; Barret et al., 2005). The genes underlying the associated regions were retrieved from the ‘67/3’ eggplant reference genome (ver. 3) (Barchi et al., 2019b). Genes were considered as potential candidates in controlling the traits assessed when carrying homozygous allelic variants classified as “high impact” according to SnpEff software v 4.2 prediction (Cingolani et al., 2012) of the eight MAGIC founders (Gramazio et al., 2019). Integrative Genomics Viewer (IGV) tool was used for visual exploration of founder genome sequences to validate SnpEff results and confirm the presence of the so-called “high impact” variants (Robinson et al., 2020). In addition, eggplant and tomato synteny was assessed by a BLASTx search of candidate genes sequences against the tomato genome (version SL4.0) in the Sol Genomics Network database (http://www.solgenomics.net).

## Author Contributions

SV, PG and JP conceived the idea and supervised the manuscript; GM, AA, MP and PG performed the field trials. All authors analyzed the results. GM and AA prepared a first draft of the manuscript and the rest of authors reviewed and edited the manuscript. All authors have read and agreed to the published version of the manuscript.

## Acknowledgements

This work was supported by the Ministerio de Ciencia, Innovación y Universidades, Agencia Estatal de Investigación and Fondo Europeo de Desarrollo Regional (grant RTI2018-094592-B-I00 from MCIU/AEI/ FEDER, UE) and European Union’s Horizon 2020 Research and Innovation Programme under grant agreement No. 677379 (G2P-SOL project: Linking genetic resources, genomes and phenotypes of Solanaceous crops). Andrea Arrones is grateful to Spanish Ministerio de Ciencia, Innovación y Universidades for a pre-doctoral (FPU18/01742) contract. Mariola Plazas is grateful to Spanish Ministerio de Ciencia e Innovación for a post-doctoral grant (IJC2019-039091-I/AEI/10.13039/501100011033). Pietro Gramazio is grateful to Spanish Ministerio de Ciencia e Innovación for a post-doctoral grant (FJC2019-038921-I/AEI/10.13039/501100011033).

## Conflict of interest

The authors have no conflict of interest to declare.

## Supporting Information

**Figure S1**. (a) Chain pollination scheme of the four-way hybrids followed to obtain the eight-way hybrids. (b) For each S0 progeny, four plants were germinated, selecting for the next generation (S1) only the first two that set fruits with viable seed. For subsequent generations, two plants were germinated and only the first one that set fruit was selected for the next generation.

**Figure S2**. Local Manhattan plot (top) and LD heatmap (bottom) surrounding the peaks PA9 (a), FA9 (b), PUC3.1 (c), PUC3.2 (d), PUC10.1 (e) and PUC10.4 (f). The red and green horizontal lines represent, respectively, FDR and Bonferroni significance thresholds. Pairwise LD between SNPs is indicated as values of *R*^*2*^ values: red indicates a value of 1 and white indicates 0.

**Figure S3**. Founder haplotype blocks representation predicted for each of the S3 individuals for the three anthocyanin-related candidate gene regions: (a) *MYB113* (SMEL_001g120500.1) on chromosome 1 between 5.11 - 5.88 Mb identified by the FA1 and PUC1 associations; (b) *COP1* (SMEL_010g339180.1.01) on chromosome 10 between 3.94 - 4.34 Mb identified by the PUC10.2 association; and (c) *MYB113* (SMEL_010g351850.1) on chromosome 10 between 91.08 - 94.81 Mb identified by the PA10, FA10 and PUC10.3 associations.

**Figure S4**. Phenotyping of the S3MEGGIC population for presence or absence of plant anthocyanins (PA), fruit anthocyanins (FA) and pigmentation under the calyx (PUC).

**Table S1**. List of candidate genes for plant anthocyanins (PA), fruit anthocyanins (FA) and anthocyanin pigmentation under the calyx (PUC).

## Notes

### Competing Interest Statement

The authors have declared no competing interest.

